## Supplementary Table 1 and 2 for "NHR-8 regulated P-glycoproteins uncouple xenobiotic stress resistance from longevity in chemosensory *C. elegans* mutants" (from another version of this preprint: version 2, 2020-09-29)

### Supplementary Table 1: Demographic Lifespan Data

| Strain | Treatment | Mean LS (days) | Median LS (days) | Mean difference from cntl | Animals (used/ total) | Max LS | Max difference from cntl | P Value | Reference cntl | Figure |
|---|---|---|---|---|---|---|---|---|---|---|
| N2 | | 23.96 | 24 | | 86/107 | 40 | | | | |
| *osm-3(dh441)IV* | | 29.54 | 33 | 23% | 62/79 | 47 | 18% | 0.0000 | N2 | 1C |
| N2 | | 24.03 | 24 | | 56/65 | 36 | | | | |
| *osm-3(dh441)IV* | | 29.35 | 31 | 18% | 65/79 | 43 | 16% | 0.0000 | N2 | |
| N2 | | 21.66 | 23 | | 92/97 | 34 | | | | |
| *osm-3(dh441)IV* | | 26.64 | 27 | 23% | 70/94 | 41 | 21%% | 0.0000 | N2 | |
| N2 | | 21.08 | 22 | | 128/145 | 33 | | | | |
| *daf-16(mu86)I* | | 17.51 | 17 | -17% | 145/153 | 26 | -21% | 0.0000 | N2 | |
| *osm-3(dh441)IV* | | 23.17 | 22 | 10% | 96/140 | 38 | 15% | 0.0014 | N2 | |
| *osm-3(dh441)IV; daf-16(mu86)I* | | 17.58 | 19 | -17% | 135/152 | 29 | -12% | 0.0000 | N2 | |
| N2 | | 21.51 | 22 | | 69/83 | 36 | | | | |
| *daf-16(mu86)I* | | 16.01 | 15 | -26% | 79/95 | 27 | -25% | 0.0000 | N2 | |
| *osm-3(dh441)IV* | | 24.80 | 24 | 15% | 69/84 | 43 | 19% | 0.0025 | N2 | |
| *osm-3(dh441)IV; daf-16(mu86)I* | | 16.24 | 15 | -25% | 83/93 | 24 | -33% | 0.0000 | N2 | 3A |
| N2 | | 20.53 | 22 | | 108/113 | 32 | | | | |
| *osm-3(dh441)IV* | | 25.53 | 26 | 24% | 91/106 | 39 | 22% | 0.0000 | N2 | |
| *nhr-8(hd117)IV* | | 16.34 | 15 | -20% | 81/93 | 26 | -19% | 0.0000 | N2 | |
| *osm-3(dh441)IV; nhr-8(hd117)IV* | | 18.27 | 15 | -11% | 88/90 | 32 | 0% | 0.0597 | N2 | |
| N2 | | 20.78 | 22 | | 105/125 | 31 | | | | |
| *osm-3(dh441)IV* | | 23.10 | 24 | 12% | 87/125 | 35 | 13% | 0.0012 | N2 | |
| *nhr-8(hd117)IV* | | 14.47 | 14 | -30% | 68/125 | 24 | -23% | 0.0000 | N2 | |
| *osm-3(dh441)IV; nhr-8(hd117)IV* | | 17.47 | 19 | -16% | 76/125 | 29 | 6% | 0.0000 | N2 | S5A |
| N2 | DMSO | 20.60 | 21 | | 89/104 | 32 | | | | |
| *osm-3(dh441)IV* | DMSO | 27.63 | 29 | 34% | 47/77 | 47 | 47% | 0.0000 | N2 | |
| N2 | Verapamil | 22.26 | 21 | 8% | 68/101 | 32 | 0% | 0.0321 | N2-DMSO | |
| *osm-3(dh441)IV* | Verapamil | 23.84 | 24 | 16%% | 45/63 | 47 | 47% | 0.0015 | N2-DMSO | |
| N2 | DMSO | 19.23 | 18 | | 127/138 | 29 | | | | |
| *osm-3(dh441)IV* | DMSO | 23.92 | 23 | 24% | 93/126 | 47 | 62% | 0.0000 | N2-DMSO | |
| N2 | Verapamil | 18.65 | 18 | -3% | 80/103 | 26 | -10% | 0.4451 | N2-DMSO | |
| *osm-3(dh441)IV* | Verapamil | 22.11 | 21 | 24% | 84/99 | 43 | 62% | 0.0002 | N2-DMSO | 4D |

Supplementary Table 2: RT-qPCR Primer Sequences

| Gene | Forward Primer Sequence | Reverse Primer Sequence |
|---|---|---|
| *act-1* | CTACGAACTTCCTGACGGACAAG | CCGGCGGACTCCATACC |
| *hsp-4* | GTGGCAAACGCGTACTGTGATGA | CGCAACGTATGATGGAGTGATTCT |
| *ero-1* | ATCGCCGCAAAGAGTGTGGAAG | TTGTCGAATTTGGGATCCGGTTTG |
| *crt-1* | TGTGGCAGGTCAAGTCAGGAAC | TTTCTTCGTCGGCCTTCTCCTTC |
| *cnx-1* | TACAATGGAGAGTGGAGCATCGG | ATTGCATGATGGCGTGCCTTTG |
| *pdi-1* | TGAGCAACTCGAGGAATTGAAGGG | AGCATCAGATTTCGCGTCCTTG |
| T14G8.3 | GGATCAAGCTAATGAGCAGCAGAC | TGACAGGTTTACAGTTCCGATGC |
| *dnj-27* | GAGAGCTTGCAGAGGTTATTGGAG | GCATCATTGGCAAGCATCCATACG |
| T24H7.2.1 | AGAGCCGAAGCTTTGTTTGAGC | ACTGATGGTGCTTGCAATTGATCC |
| *xbp-1u* | AGAAGTCGTCGGTGAGGTTG | CCTGTTCCCACTGCTGAG |
| *xbp-1s* | TGCCTTTGAATCAGCAGTGG | ACCGTCTGCTCCTTCCTCAATG |
| *pgp-1* | TGGCTCCGAGATCAAGACGTTG | TGAGCCATGGTAACTGAGGAAGG |
| *pgp-2* | AGGAGGTTGAGGAATGCAAGGC | CCATCCGACTTTGTCTCCGTTG |
| *pgp-3* | TTGCTAGAGCGATTGTCAGAGACC | TGTCAAGAGCTGACGTGGCTTC |
| *pgp-4* | AAGATGACACCACCCAAGATGATG | TGTCGAAAGACGTCGGGAGATTG |
| *pgp-5* | ATCCGGTCTGTTGGACTCTTGG | TACCGTTCAACTTCTTGCTGCTG |
| *pgp-6* | GAAACAGCGGTCGTTGTTAGGC | CAGTGCTTCACTAAGACAGAAGGC |
| *pgp-7* | TGCTGCTACCGAAACGTCTATCC | CCAACGCTTCAATCGCAGTTC |
| *pgp-8* | AAGCATCAGAAGGGCGGACAAC | CGCAATGACGAATAGTGCTAAGCC |
| *pgp-9* | CGTGTATTGTAGTGGCTCATCGG | TC A GTTC A TTGTGGGTTC C TTGC |
| *pgp-10* | TAAGAGAACCACCCGTTGACATGG | AGAGGTTTCCTTCCAGCTTCGG |
| *pgp-11* | CACTTCGATCGGCAACAACTGG | TTGAACAGTACTCAGTCGATGTGC |
| *pgp-12* | ATGCGATCGAGCCAATTGATGG | CCTCAGTCGACACATTCACAGTTG |
| *pgp-13* | CGAAGCTGTCAACTACTGCATTTC | GCATCATCGCCATGAGACATCTG |
| *pgp-14* | TGCTTCTGGATGAAGCCACGTCAG | AATGCCTCTTGTACAGCCCTTTCG |
